# TLN468 changes the pattern of tRNA used to read through premature termination codons in CFTR

**DOI:** 10.1101/2023.02.02.526440

**Authors:** Sabrina Karri, David Cornu, Claudia Serot, Lynda Biri, Aurélie Hatton, Iwona Pranke, Isabelle Sermet-Gaudelus, Alexandre Hinzpeter, Laure Bidou, Olivier Namy

## Abstract

Nonsense mutations account for 12% of cystic fibrosis (CF) cases. The presence of a premature termination codon (PTC) leads to gene inactivation, which can be countered by the use of drugs stimulating PTC readthrough, restoring production of the full-length protein. We recently identified a new readthrough inducer, TLN468, more efficient than gentamicin.

We measured the readthrough induced by these two drugs with different cystic fibrosis transmembrane conductance regulator (*CFTR*) PTCs. We then determined the amino acids inserted at the S1196X, G542X, W846X and E1417X PTCs of *CFTR* during readthrough induced by gentamicin or TLN468. TLN468 significantly promoted the incorporation of one specific amino acid, whereas gentamicin did not greatly modify the proportions of the various amino acids incorporated relative to basal conditions. The function of the engineered missense CFTR channels corresponding to these four PTCs was assessed with and without potentiator. For the recoded CFTR, except for E1417Q and G542W, the PTC readthrough induced by TLN468 allowed the expression of CFTR variants that were correctly processed and had significant activity that was enhanced by CFTR modulators. These results suggest that it would be relevant to assess the therapeutic benefit of TLN468 PTC suppression in combination with CFTR modulators in preclinical assays.

## INTRODUCTION

Cystic fibrosis (CF) is an autosomal recessive disease caused by mutations of the gene encoding the cystic fibrosis transmembrane conductance regulator (CFTR) protein. Treatments, including CFTR modulators, have proven clinical benefits for patients carrying the F508del mutation (1). However, such treatments are ineffective in patients carrying nonsense mutations introducing a premature termination codon (PTC). These patients account for about 12% of all CF patients in Europe (2).

The introduction of an in-frame stop codon (PTC) has two major deleterious effects. First, PTC-containing mRNAs are frequently unstable, resulting in a large decrease in steady-state mRNA abundance. Second, the PTC leads to the synthesis of truncated proteins that are often non-functional. Together, these two PTC-induced events substantially decrease functional protein levels, potentially resulting in severe disease. Therapeutic approaches are being developed to promote the translational readthrough of the PTC, favoring the recruitment of a near-cognate tRNA in place of the termination complex, thereby enabling the production of a full-length protein (3).

Over the last decade, much research has focused on in-frame PTCs as potential therapeutic targets. The best-characterized drugs active against PTCs are aminoglycosides. Their readthrough efficiency has been evaluated for several genes, including the cystic fibrosis transmembrane conductance regulator (CFTR) gene (4). Encouraging results have been obtained in some cases, particularly for mutations displaying high levels of readthrough in the presence of gentamicin (5). However, despite their medical value, aminoglycosides are effective against only a small set of mutations and these drugs have adverse effects, with reports of various levels of ototoxicity and/or nephrotoxicity (6,7).

The limitations of aminoglycosides have led to the development of other compounds with different structures. The various new molecules developed include ataluren (8), negamycin (9,10), clitocine (11), escin (12), 2,6-diaminopurine (13) and, more recently, SRI-41315 (14). Further studies will be required to determine the true clinical potential of these molecules. The identification of new readthrough drugs remains important because one of the main factors limiting the impact of readthrough-inducing treatments is their highly variable efficacy (4). Indeed, treatment response is strongly influenced by the nature of the stop codon (UAG, UGA or UAA) and the nucleotide sequence surrounding it (15–17), restricting the potential clinical benefits of these drugs to a limited number of patients. Another important factor is the activity of the recoded proteins. Several different tRNAs have been shown to be incorporated by readthrough at the stop codon. If these tRNAs do not allow the insertion of the amino acid present in the WT protein, then there is a risk of obtaining a protein with little or no activity. Amino acids inserted were initially identified by mass spectrometry in yeast (18) and have more recently been identified in mammalian expression systems (19). However, there have been no systematic studies of the identity of the amino acids incorporated during treatment with readthrough inducers.

We recently identified a new readthrough inducer, TLN468 (20), with a different mode of action and greater efficacy than gentamicin. We characterized the effect of this new drug on PTCs identified in CF patients. We first identified TLN468-induced readthrough levels for 12 of the most frequent *CFTR* PTCs introduced by nonsense mutations of the CFTR gene. We then selected four PTCs for further studies based on their ability to respond to TLN468 and to represent the three different stop codons: S1196X (UGA), G542X (UGA), W846X (UAG) and E1417X (UAA). For each mutation, we determined basal and drug-induced readthrough levels, the identity of the amino acids incorporated, and the activity of the restored proteins. Readthrough efficiencies were highly variable depending on the PTC, confirming that PTC suppression depends on the nature of the codon and the nucleotide context. Surprisingly, the proportions of the amino-acids introduced in the presence of TLN468 differed significantly from those introduced in the presence of gentamicin or in basal conditions. Two to five different amino acids were incorporated in basal conditions, and the proportions of these amino acids were modified moderately by gentamicin treatment. By contrast, TLN468 promoted the almost exclusive incorporation of cysteine for the UGA codon, glutamine for the UAG codon and tyrosine for the UAA codon. This new readthrough inducer thus displays a specificity that must be taken into account for therapeutic purposes.

We investigated whether CFTR proteins with these amino acids in place of the PTC would be functional, by engineering missense substitutions at these positions and assessing the activity of the resulting protein. We found that all CFTR variants other than G542C and G542W were appropriately processed and active, demonstrating a high potential for nonsense suppression therapy. Moreover, the CFTR potentiator VX-770 (Ivacaftor) (21) increased the activity of these proteins, thereby increasing their potential therapeutic benefits. These results suggest that combining the new nonsense suppression drug, TLN468, with CFTR modulators is a promising approach for the development of therapies maximizing CFTR function in CF patients with nonsense mutations.

## Results

We first measured basal and TLN468-induced readthrough levels associated with 12 *CFTR* PTCs (listed in Table 1) in human HeLa cells (Figure 1). We assessed the ability of TLN468 to stimulate readthrough by using a dual reporter system to quantify PTC readthrough efficiency (22). This reporter system includes two enzymatic activities (β-galactosidase and firefly luciferase), β-galactosidase being used as an internal control for the normalization of expression levels. The local mRNA sequence context flanking a PTC is known to be determinant for readthrough (23). We therefore introduced the PTC within its nucleotide context (nine nucleotides on either side of the PTC) in this reporter construct, at the start of the luciferase coding phase. For each molecule tested, we performed at least four independent transfections and measurements (Figure 1).

**Figure 1:**
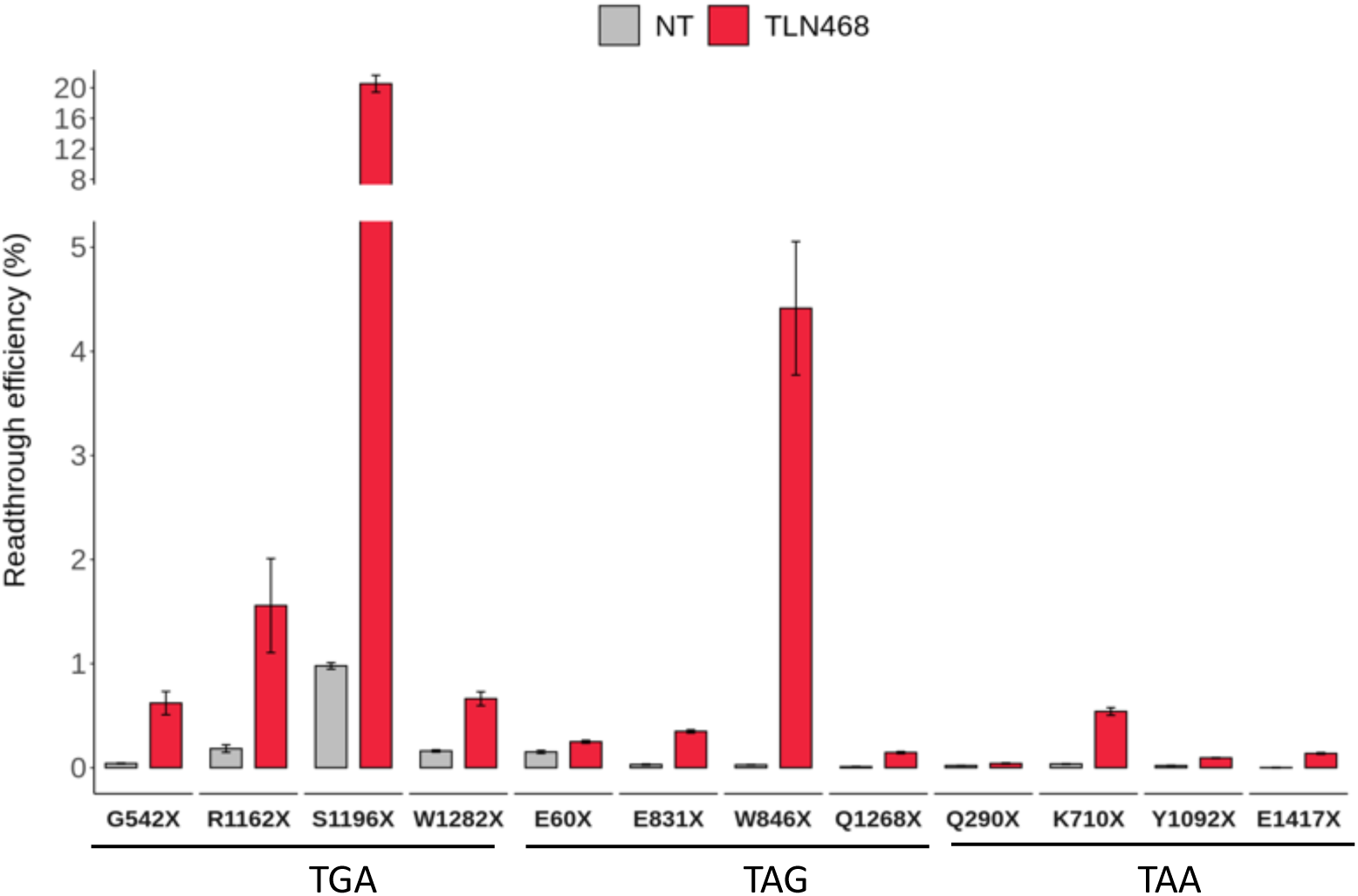
Quantification of the PTC readthrough efficiency. Readthrough levels measured for the 12 CFTR PTCs listed in table 1, in the basal state (NT) and after incubation with 80 µM TLN468 for 24 h in the Expi 293F human cell line. The y-axis was interrupted above 5% and the scale was modified to make it possible to indicate the very high % readthrough of S1196X without obliterating the other values. The plot shows the mean and standard deviation, n>4 for all samples. For all mutation, difference between non treated (NT) and TLN468-induced readthrough is significant (p<0.0008 unpaired t-test). N=5 for all samples except the WT (n=8 for basal conditions and n=10 with VX-770).

**Table 1:**
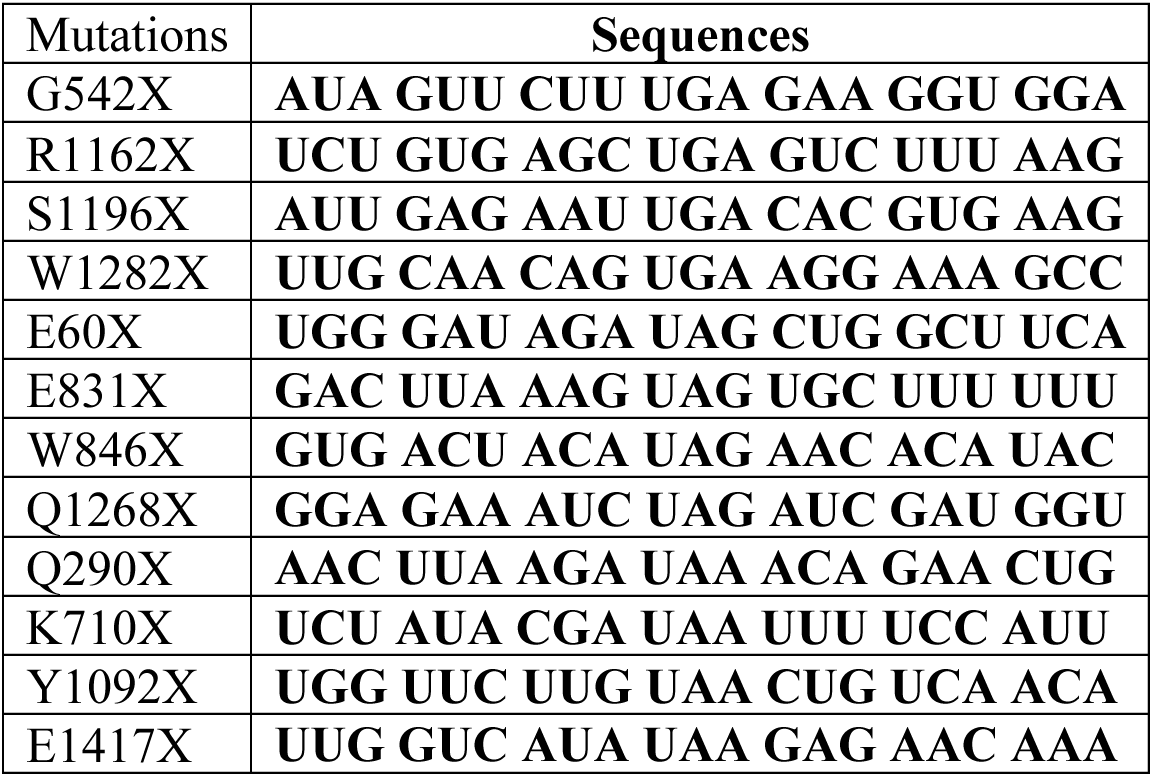
Selected nonsense mutations. Four sequences were selected for each stop codon and inserted, together with their flanking sequences, into the dual reporter plasmid.

Basal readthrough was highly variable, ranging from 0.002% for E1417X to 1% for S1196X. TLN468 was effective for all stop mutations and stimulated readthrough to levels ranging from 0.14% for E1417X (increase by a factor of 70) to 20% for S1196X (increase by a factor of 20).

Four PTCs were then selected for further studies based on their ability to respond to TLN468 and to represent the three different stop codons: S1196X (UGA), G542X (UGA), W846X (UAG) and E1417X (UAA). We compared the effects on readthrough of gentamicin and TLN468 for these four mutations (Figure S1).

Interestingly, except for G542X, TLN468 induced readthrough much more efficiently than gentamicin, for all three stop codons. TLN468 was twice as effective as gentamicin for S1196X (UGA) and E1417X (UAA) and more than 12 times more effective than gentamicin for W846X, increasing the rate of readthrough from about 0.3% with gentamicin to 4.4% with TLN468.

### Identification of the amino acids incorporated during translational suppression of the *CFTR* S1196X, G542X, W846X and E1417X nonsense mutations

We investigated the identity of the natural tRNA suppressors of each PTC (Figure 2). PTC suppression occurs at relatively low frequencies, making it difficult to analyze the amino acids inserted during this process. We overcame this difficulty by using a recently developed GST reporter allowing the efficient single-step purification of the protein synthesized during a stop codon-readthrough event (18). The GST coding sequence of this reporter has been humanized to enhance its expression in human cells. Moreover, based on an N-terminal protein degradation frequently observed in cells, we introduced a spacer region immediately downstream from the AUG of GST but upstream from the insertion site of the PTC sequences. GST is produced only if ribosomes read through the inserted stop codon.

**Figure 2:**
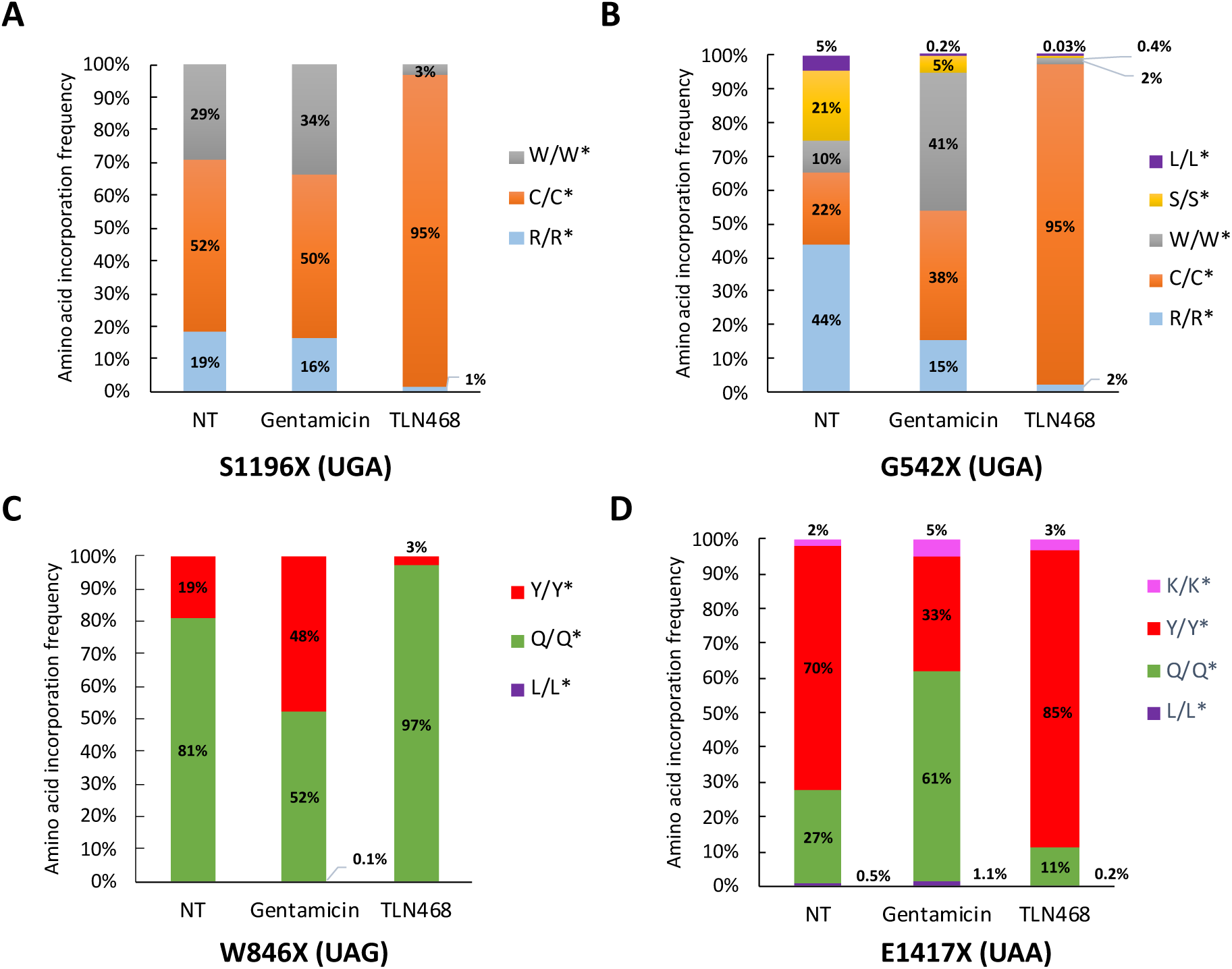
Relative frequencies of the readthrough amino acids incorporated at CFTR stop mutations in the Expi293F human cell line, as established by MS-MS. Comparison of the amino acids inserted in the basal state (NT) and following gentamicin (2.5 mM)- or TLN468 (80 µM)-induced suppression of the S1196X (A), G542X (B), W846X (C) and E1417X (D) PTCs. X/X*: ratio of the peptide purified from human cells (X) to the corresponding AQUA peptide (X*). n=3 sample points.

We identified the amino acids inserted during basal readthrough of the *CFTR* stop mutations or after PTC suppression induced by gentamicin or TLN468 treatment. The readthrough GST proteins were purified and the peptides obtained were analyzed by LC-MS/MS with systematic normalization against the corresponding synthetic peptide.

In basal conditions, three amino acids were inserted, with different frequencies, at the *CFTR* S1196X UGA codon: cysteine predominated, but arginine and tryptophan were also observed (Figure 2.A). The same three amino acids were inserted at the *CFTR* G542X UGA codon, but with arginine predominating in this context. Surprisingly, we also observed the incorporation of serine (21%) and leucine (4.6%) (Figure 2.B). In parallel, at the *CFTR* W846X UAG and E1417X UAA codons, we observed the insertion of glutamine, which predominated for the UAG codon, and tyrosine, which predominated for the UAA codon (Figure 2.C and D). We also found a small portion of lysine (1.7%) and leucine (0.5%) insertions at the E1417X UAA stop codon (Figure 2.D). Interestingly, gentamicin modified the proportions of the various amino acids inserted according to the mutation. For S1196X, gentamicin had no significant effect on the proportions of the three amino acids incorporated, but for G542X, it favored the incorporation of tryptophan and cysteine at the expense of the other three amino acids. For W846X and E1417X, glutamine and tyrosine remained the predominant amino acids inserted after gentamicin treatment, but with a modification of their respective proportions. TLN468 had a more profound impact on tRNA incorporation, as it promoted the incorporation of a single amino acid, the identity of which depended on the PTC (>95%) for each of the mutations tested. For the UGA PTCs (Figure 2.A & 2.B), TLN468 promoted the incorporation of cysteine (95%). For the UAG codon, it promoted the incorporation of glutamine in 97% of cases (Figure 2.C) and for the UAA codon, it promoted the incorporation of tyrosine in 85% of cases (Figure 2.D).

### Lowering the dose of TLN468 does not affect amino-acid incorporation frequency

We then investigated whether the efficiency of amino-acid incorporation varied as a function of the TLN468 concentration used. We addressed this question using S1196X, which had the highest readthrough efficiency following treatment with TLN468 at the maximum dose of 80 μM. We first determined the readthrough efficiency of TLN468 at various doses (0, 20, 40, 60 and 80 μM) with the S1196X stop codon. As expected, effect of TLN468 was dose-dependent, with the highest readthrough efficiency at 80 μM (20%), decreasing to 15.9% at 60 μM, 8.7% at 40 μM and 5% at 20 μM (Figure 3.A).

**Figure 3:**
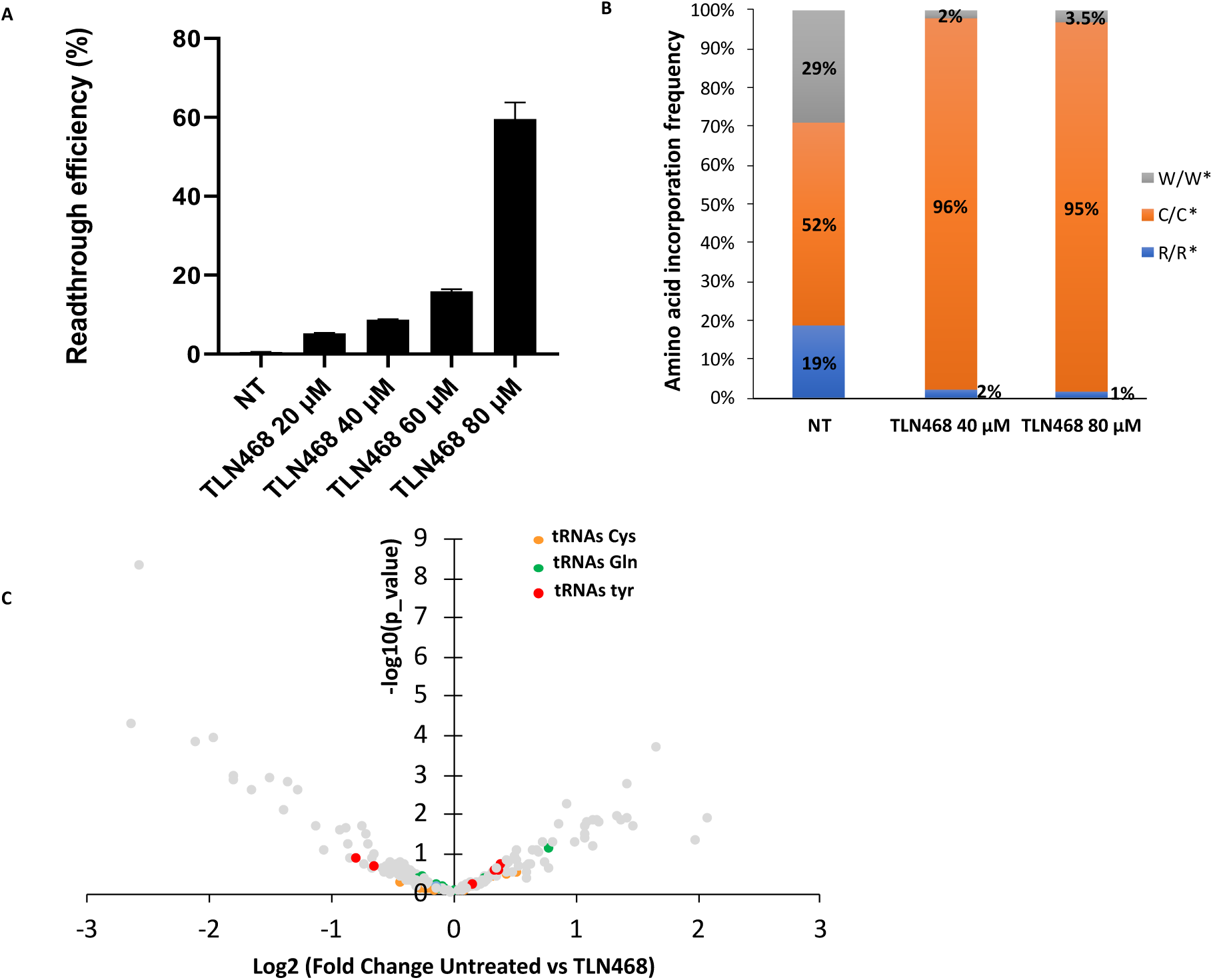
Dose effect of TLN468 on readthrough. (A) Quantification of readthrough efficiency at the CFTR S1196X PTC with various concentrations of TLN468 in Expi 293F human cells. The plot shows the mean and standard deviation, n=5, for all samples. (B) Comparison of the amino acids inserted during TLN468-induced suppression of the S1196X PTC at 40 µM and 80 µM. X/X*: ratio of the peptide purified from human cells (X) to the corresponding AQUA peptide (X*). n=3 sample points. (C) Impact of TLN468 on the tRNA population. Volcano plot of variations in the pool of tRNA isodecoders between untreated Expi293F cells and Expi293F cells treated with TLN468 (80 µM). The volcano plot was generated with the CPM (counts per million reads mapped) values calculated with multimapped reads. n=3 samples.

We then identified the amino acids inserted following treatment with 80 µM and 40 µM TLN468. We observed that cysteine was systematically the main amino acid incorporated at the S1196X stop codon (Figure 3B). These findings suggest that, regardless of the dose used, TLN468 acts strongly on the cysteine tRNA, enabling it to be incorporated more efficiently.

We checked that this effect was not due to the overexpression of cysteine tRNA isodecoders in response to TLN468 treatment, by performing a tRNA-seq experiment. The quantification of all tRNA isodecoders in Expi293 cells in the presence and absence of TLN468 revealed no difference in cysteine tRNA abundance (Figure 3C), ruling out this possibility. Similarly, glutamine and tyrosine tRNAs, which were predominantly incorporated at UAG and UAA stop codons, respectively, after TLN468 treatment, were not overexpressed either.

### Maturation and function of the predicted recoded channels

The level of activity of CFTR variants may depend on the amino acid incorporated. We investigated the effect on protein maturation and function of the amino acids incorporated in the presence of TLN468 for the *CFTR* S1196X, G542X, W846X and E1417X mutations. Based on our knowledge of the amino acids incorporated at these positions, we were able to mutate a WT *CFTR* cDNA at these four positions to create twelve CFTR variants with different sense codons corresponding to the amino acids identified by mass spectrometry (Figure 2).

We transfected HEK293 cells with these CFTR variants and assessed the production and maturation of the CFTR protein. We explored CFTR maturation, by calculating the ratio of CFTR band C (the mature, fully glycosylated form) to CFTR band B (the immature, non-glycosylated form) on western blots (Figure 4.A). Most of the WT CFTR was present as Band C, indicating that, in WT conditions, CFTR proteins are almost fully glycosylated (about 80%). The WT protein was considered to display 100% maturation. We observed slightly lower levels of glycosylation for the G542W (67% WT levels) and G542C (72% WT levels) variants (Figure 5.B). All the other variants were efficiently glycosylated to at least 85% WT CFTR levels (Figure 4.B).

**Figure 4:**
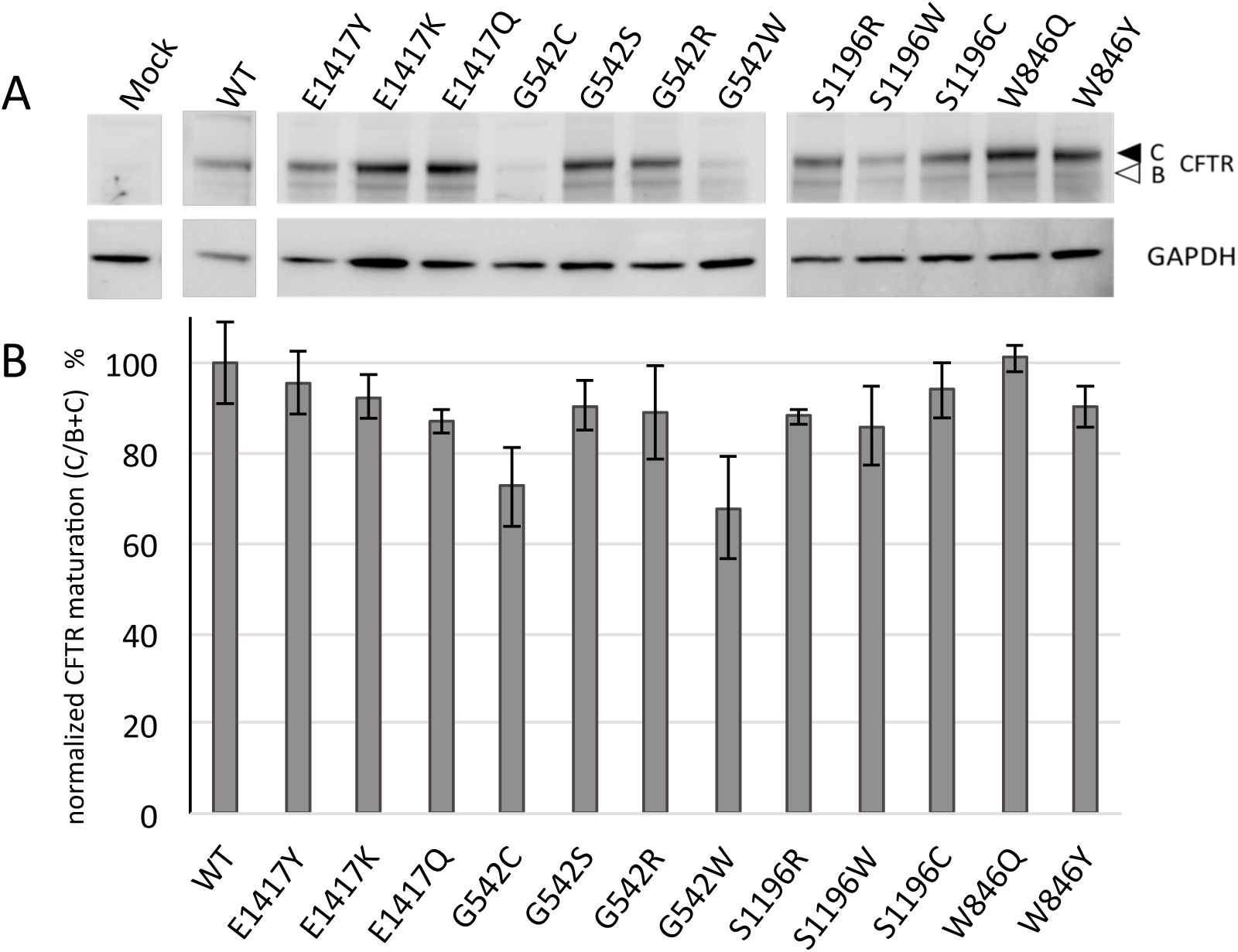
Maturation of recoded cystic fibrosis transmembrane conductance regulator (CFTR) channels. (A) Representative western blot of HEK293 cells expressing the indicated construct. CFTR bands C and B are indicated. (B) CFTR maturation was assessed by determining the C/(B+C) ratio, with the WT CFTR considered to present 100% maturation. Each band was normalized against GAPDH. n=3 (minimum) for all samples.

**Figure 5:**
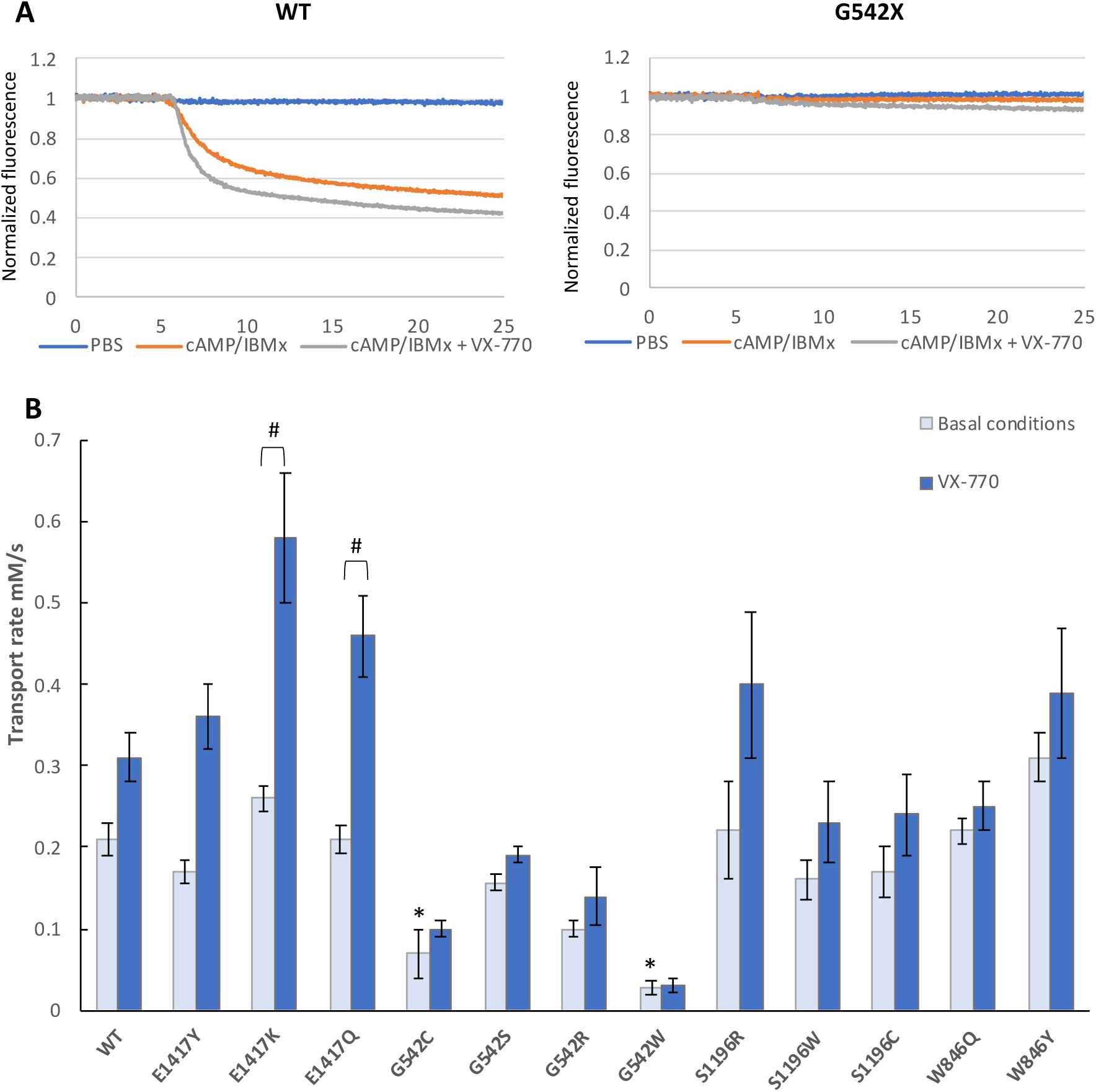
Functioning of the recoded cystic fibrosis transmembrane conductance regulator (CFTR) channels. (A) Representative recordings obtained from HEK293 cells expressing the indicated construct (CFTR-WT or G542W) after incubation with PBS or cAMP/IBMx-containing PBS, with or without the CFTR potentiator VX-770. (B) Transport rates measured under basal conditions and in the presence of the CFTR potentiator VX-770. *: significant difference compared with wild-type control (p<0.01, ANOVA); #: significant effect of VX-770 (p<0.01, unpaired t-test). N=4 for all samples except for the WT (n=8 for basal conditions and n=10 with VX-770).

The functioning of the CFTR mutants was then assessed in a halide-sensitive YFP-based assay. This assay is based on the sensitivity of the YFP protein to the I^-^ ion. In iodide-rich conditions, activated CFTR induces an influx of iodide ions that quenches YFP fluorescence. In this assay, we used a genetically modified YFP with a high quenching sensitivity to iodide but a very low sensitivity to chloride generated through the introduction of key mutations (H148Q-I152L) into the YFP protein (24).

The iodide transport rate was 0.21±0.02 mM·s^−1^ in WT-transfected cells (Figure 5B). Nine CFTR variants had activity levels similar to or slightly lower than that of the WT protein, whereas three of the four variants at position 542 (G542W, G542R, G542C) clearly displayed lower rates of transport, less than half that of the WT protein (Figure 5B). The two mutants with the lowest levels of activity, G542W and G542C, had maturation defects.

Cells were also incubated with the CFTR potentiator VX-770, which is known to improve CFTR channel gating. As expected, this molecule induced a clear increase in iodide transport rate. WT-expressing cells presented a 1.5-fold increase in iodide transport rate (0.31±0.03 mM·s^−1^; *p*<0.0001) (Figure 5B). A similar increase was observed with the proteins presenting normal levels of basal activity except for W846Y and W846Q (1.2- and 1.1-fold increases, respectively). The strongest effect of VX-770 was observed with E1417K-expressing cells, in which iodide transport rate increased by a factor of 2.2 (0.58±0.05 mM·s^−1^). For the three channels mutated at position 542, which had the lowest levels of band C (G542W, G542R and G542C), the potentiator did not significantly enhance channel function.

## Discussion

A partial restoration of protein function by PTC suppression has been demonstrated in several diseases caused by nonsense mutations, including CF (9,25,26). Moreover, PTC suppression in combination with a corrector, such as lumacaftor, has been shown to promote a further rescue of expression and function for CFTR PTC mutations (19,27,28).

We report here the first systematic analysis of the amino acids incorporated by stop codon readthrough in basal conditions and in the presence of either gentamicin or TLN468 in human cells. As expected, basal readthrough was highly variable for the 12 PTCs studied, with lowest rate obtained for a UAA (E1417X) stop codon, which has been reported to be the strongest termination codon, and the highest rate obtained for a UGA (S1196X) stop codon, which has been reported to be the leakiest stop, especially if the base immediately 3’ is a C, which is the case for the S1196X PTC (27). Treatment with TLN468, a new non-aminoglycoside readthrough inducer, is globally more effective than treatment with gentamicin for inducing readthrough. The difference between basal and drug induced readthrough rates was highest for E1417X PTC, but the highest induced rate of readthrough was that for S1196X, which also had particularly high basal readthrough levels.

The mode of action of gentamicin is well documented, but it remains unknown how TLN468 stimulates stop codon readthrough. This molecule represents a new chemical family of readthrough inducers, expanding the range of products available to suppress PTC-related diseases. An understanding of the effect of TLN468 on tRNA incorporation may provide interesting clues to the molecular target of TLN468. In this context, in addition to measuring effective protein production by quantitative methods, it is crucial to demonstrate that the protein generated by readthrough is functional. Here, we assessed the readthrough efficiency of TLN468 for 12 PTCs in the *CFTR* gene, subsequently focusing on four of these mutations to determine which amino acids were inserted in the presence of the drug: S1196X, G542X, W846X and E1417X. This is the first comprehensive study aiming to determine which amino acids are incorporated during stop codon readthrough in the presence and absence of a readthrough-inducing drug.

Interestingly, the amino acids incorporated in the absence of the drug closely resembled those previously identified in yeast (18), although a greater variety of amino acids were incorporated in humans than in yeast for the G542X mutation. Indeed, we report here the first identification of a serine residue incorporated during PTC readthrough. A comparison of the amino acids found in untreated cells at S1196X and G542X, two UAG stop codons, clearly indicated that, contrary to initial assumptions, the nucleotide environment of the PTC plays a role in tRNA selection during readthrough. We then focused on the identity of the amino acids incorporated in the presence of readthrough-inducing drugs (gentamicin or TLN468). Gentamicin had an unexpected effect, clearly increasing the incorporation of Trp (W) at G542X but not at S1196X, despite both being UGA codons. It stimulated the incorporation of tyrosine at W846X (UGA) but restricted the incorporation of this amino acid at E1417X (UAA). This is surprising because gentamicin acts by binding the decoding center and promoting the incorporation of near-cognate tRNAs (29). We might therefore expect that all the tRNAs incorporated under basal conditions would also be incorporated after gentamicin treatment, with no change in their proportions, as observed for S1196X. Unfortunately, too little is known about the effect of gentamicin on tRNAs. Indeed, previous studies analyzing amino-acid incorporation did not include untreated conditions (probably because readthrough levels were too low for quantitative analysis), but our findings suggest that the effect of gentamicin depends on a combination of factors (nucleotide context, stop codon) favoring a subgroup of tRNAs.

The mode of action of TLN468 is unknown, although a recent *in vitro* study indicated that TLN468 did not inhibit peptide release activity, suggesting that it may act by stimulating tRNAs directly, rather than inhibiting the activity of release factors (30). We found that, even at low concentrations, TLN468 strongly biased tRNA incorporation, systematically favoring the almost exclusive incorporation of a single amino acid regardless of the nucleotide context of the PTC. However, the identity of the favored amino acid differed between types of stop codon. It is currently difficult to identify a feature common to the three tRNAs for which incorporation is strongly stimulated by the presence of TLN468 (i.e Cys, Gln, Tyr). None of them is overexpressed in treated cells (Figure 3.C), excluding the possibility of a disturbance of tRNA homeostasis. It is possible that the drug alters the chemical modifications present on tRNAs. Unfortunately, in the case of human tRNAs, not all of these modifications are known, making an extensive study difficult. For the known modifications, we found no differences common to the tRNAs allowing the incorporation of cysteine, glutamine and tyrosine.

Clinically, the activity of the restored protein is a very important outcome. Our work clearly revealed the insertion of several different amino acids by readthrough at a given PTC. It was therefore important to determine whether the corresponding CFTR proteins would display any activity. Our results indicate that for the G542 position, the identity of the amino acid is crucial for CFTR activity, as reported in other studies (19). However, for the other positions tested, substitutions had no major effect on CFTR activity, particularly in the presence of VX-770, opening up the possibility that even if an amino acid different from that in the wild-type protein is incorporated, the resulting protein can display sufficient activity to have a physiological effect. In conclusion, our work demonstrates that TLN468 is a very promising candidate drug for treating cystic fibrosis caused by a nonsense mutation, through its ability to induce the targeted incorporation of a single amino acid at the position of the PTC.

## MATERIALS AND METHODS

### Cell lines and media

Expi293F cells (Gibco), human cells derived from suspension cultures of HEK293 cells, were used to measure readthrough efficiency and the rates of incorporation of the various amino acids. They can grow to high density in Expi293™ Expression Medium (Gibco) with queuine added directly to the medium at a final concentration of 10 nM. Cells were transfected in the presence of Expifectamine (Gibco).

HeLa cells were used to measure readthrough efficiency and to determine the relative frequencies of incorporation for the various amino acids. These cells were cultured in DMEM supplemented with 10% fetal calf serum and were transfected in the presence of jetPEI (Invitrogen).

HEK293 cells were used for western blots and functional assays. These cells were cultured in DMEM supplemented with 10% fetal calf serum and were transfected in the presence of Lipofectamine 2000 (Life Technologies).

### Readthrough quantification

For each nonsense mutation tested, complementary oligonucleotides corresponding to the stop codon and nine nucleotides on either side of the stop codon were ligated into the pAC99 dual reporter plasmid, as previously described (22). This dual reporter plasmid was used to quantify stop codon readthrough through the measurement of luciferase and beta galactosidase (internal normalization) activities, as previously described (31). Readthrough levels for nonsense mutations were analyzed in the presence or absence of the tested molecules. The cells were used to seed a six-well plate. The following day, they were transfected with the reporter plasmid in the presence of jetPEI reagent (Invitrogen). They were incubated for 14 h and then rinsed with fresh medium, with or without potential readthrough inducers. Cells were harvested 24 hours later with trypsin–EDTA (Invitrogen), and beta-galactosidase and luciferase activities were assayed as previously described (31). Readthrough efficiency was estimated by calculating the ratio of luciferase activity to beta-galactosidase activity obtained with the test construct, with normalization against an in-frame control construct.

### Purification of readthrough proteins

Readthrough proteins were produced with pCDNA3.4-GST. This vector was constructed by adapting the codon frequencies of the GST gene from pYX24-GST to match human values and adding a 171nt (H4) linker sequence (32) at the AfeI site immediately downstream from the start codon.

Expi293F cells were transfected with pCDNA3.4-GST to express a readthrough GST protein. The drug was added to the medium the following day. Cell extracts were obtained 24 hours after treatment and GST was purified on a 1 mL GSTrap HP column (GE Healthcare) with an AKTA purifier 10 FPLC system, as previously described (18). The purification product was subjected to SDS-PAGE to separate the *S. japonicum* GST from endogenous GST for mass spectrometry .

### Mass spectrometry

Mass spectrometry analysis was performed on Coomassie blue-stained gel bands corresponding to the readthrough GST proteins (18). Briefly, the bands were subjected to in-gel LysC digestion as previously described and the readthrough peptides were further identified and quantified by nanoLC–MSMS with a nanoElute liquid chromatography system (Bruker) coupled to a timsTOF Pro mass spectrometer (Bruker). Before MS analysis, peptides were cleaned with ZipTip C18 (Millipore), dried under vacuum and then resuspended in 0.1% n-octylglucopyranoside (Sigma–Aldrich) to prevent the loss of hydrophobic peptides. Synthetic heavy isotope-labeled peptides (AQUA, Thermo) were used to spike the mixture of their endogenous counterparts to make it possible to take the relative ionization efficiencies of peptides and matrix effects into account. Peptides were loaded in solvent A and separated on an Aurora analytical column (ION OPTIK, 25 cm × 75 m, C18, 1.6 m) with a gradient of 0– 35% solvent B over 30 minutes. Solvent A was 0.1% formic acid and 2% acetonitrile in water and solvent B was 0.1% formic acid in acetonitrile. MS and MS/MS spectra were recorded from m/z 100 to 1700 with a mobility scan range from 0.6 to 1.4 V s/cm^2^. MS /MS spectra were acquired in PASEF (parallel accumulation serial fragmentation) ion mobility-based acquisition mode with the number of PASEF MS/MS scans set to 10. Raw data for MS and MS/MS were processed and converted into mgf files with DaUAAnalysis software (Bruker). Peptides and proteins were identified by using the MASCOT search engine (Matrix Science, London, UK) to query a homemade database (the unknown amino acid at the stop codon is represented by “X” in the readthrough GST sequence). Database searches were based on LysC cleavage specificity with two possible missed cleavages. The carbamidomethylation of cysteines was set as a fixed modification and the oxidations of methionine and tryptophan were considered as variable modifications. Peptide and fragment tolerances were set at 10 ppm and 0.05 Da, respectively. Only ions with a score higher than the identity threshold and a false-positive discovery rate of less than 1% (Mascot decoy option) were considered.

The XIC-based relative quantification of readthrough peptides was corrected by dividing the XIC peak areas for endogenous peptides by those of their heavy isotope-labeled counterparts.

### Mutagenesis

All mutants were generated by site-directed mutagenesis with the QuikChange XL II site-directed mutagenesis kit (Agilent, Santa Clara, CA, USA), as previously described (33).

### Western blotting

HEK293 cells were transfected with the appropriate constructs and harvested after 48 h. Cells were lysed with Ripa Buffer (Thermo Fisher Scientific) supplemented with 1% protease inhibitor cocktail (Roche) and incubated on ice for 30 minutes. Total protein was quantified with Bradford reagent (Biorad) and samples containing 30 µg total protein were analyzed by western blotting on 4-12% Bis-Tris Gels (Invitrogen). Proteins were transferred onto nitrocellulose membranes in accordance with the manufacturer’s instructions (Biorad). Membranes were saturated by overnight incubation in 5% skim milk powder, 5% BSA in phosphate-buffered saline (PBS) and incubated for 1 h with the 596 anti-CFTR antibody (1/1000, CFFoundation). Membranes were washed three times in 0.1% Tween in PBS and then incubated with the secondary antibody (horseradish peroxidase–conjugated anti-mouse IgG [1/2,500]) for 1 h. The membranes were washed five times and chemiluminescence was detected with ECL Prime Western Blotting Detection Reagents (Amersham, GE Healthcare). The signal was quantified with ImageLab software.

### YFP-based functional assay

CFTR activity was measured in transiently transfected HEK293 cells with the halide-sensitive yellow fluorescent protein YFP-H148Q/I152L (kindly provided by L.J. Galietta; Telethon Institute of Genetics and Medicine (TIGEM), Naples, Italy). Transfected cells were incubated for 30 min with 100 μL of PBS or PBS supplemented with CPT-cAMP, a cell-permeant cAMP analog required for CFTR activation (100 μM; Sigma Aldrich), and IBMX, which potentiates cAMP-dependent CFTR activity (100 µM) in the presence or absence of VX-770 (10 μM; Selleckchem). Cell fluorescence was measured continuously with a CLARIOStar (BMG Labtech) before and after the addition of 200 μL PBS-NaI. The resulting curves were then normalized against the initial fluorescence and an exponential function was fitted to the signal decay for derivation of the maximum slope. Maximal slopes were converted to rates of change in intracellular iodide concentration (in mM·s^−1^).

### tRNA sequencing

Total RNA was extracted from Expi293F cells with and without treatment for 24 h with 80 µM TLN468. Extractions were performed in triplicate. tRNAs were extracted by gel purification from 5 μg total RNA, in triplicate, and were sequenced by ArrayStar Inc. (USA). ArrayStar Inc. also performed the bioinformatics analysis, by aligning tRNA reads with high confidence tRNA sequences from GtRNAdb. The tRNAseq data have been deposited in the GEO database.

### Statistics

Quantitative variables are expressed as the mean±SD. Comparisons to wild-type (WT) conditions were made by one-way ANOVA followed by Fisher’s tests. In CFTR treatment analysis, unpaired *t*-tests were used to calculate the significance of differences.

## Author contributions

SK, CS and LB performed readthrough and western-blot experiments; SK and DC performed mass spectrometry analysis; SK, LB, SC, BC and AH performed mutagenesis and functional experiments, LB and ON designed the experiments and analyzed the results. All the authors participated in the writing of the manuscript.

## Acknowledgments

We thank the proteomics facility of I2BC for mass-spectrometry expertise and analysis. This work was funded by grants awarded to ON by *Vaincre la Mucoviscidose* (No. RF20180502275), the ANR (grant Actimeth (19-CE12-0004-02)) and AFM-Telethon (grant 19660 and translectin No. 20531). SK is supported by a *Vaincre la Mucoviscidose* grant awarded to ON (No. RF20180502275). *Vaincre la Mucoviscidose* played no role in the design of the study, or the analysis or interpretation of the data. English has been edited by Alex Edelman & associates.

## Supplementary data

**Figure S1:**
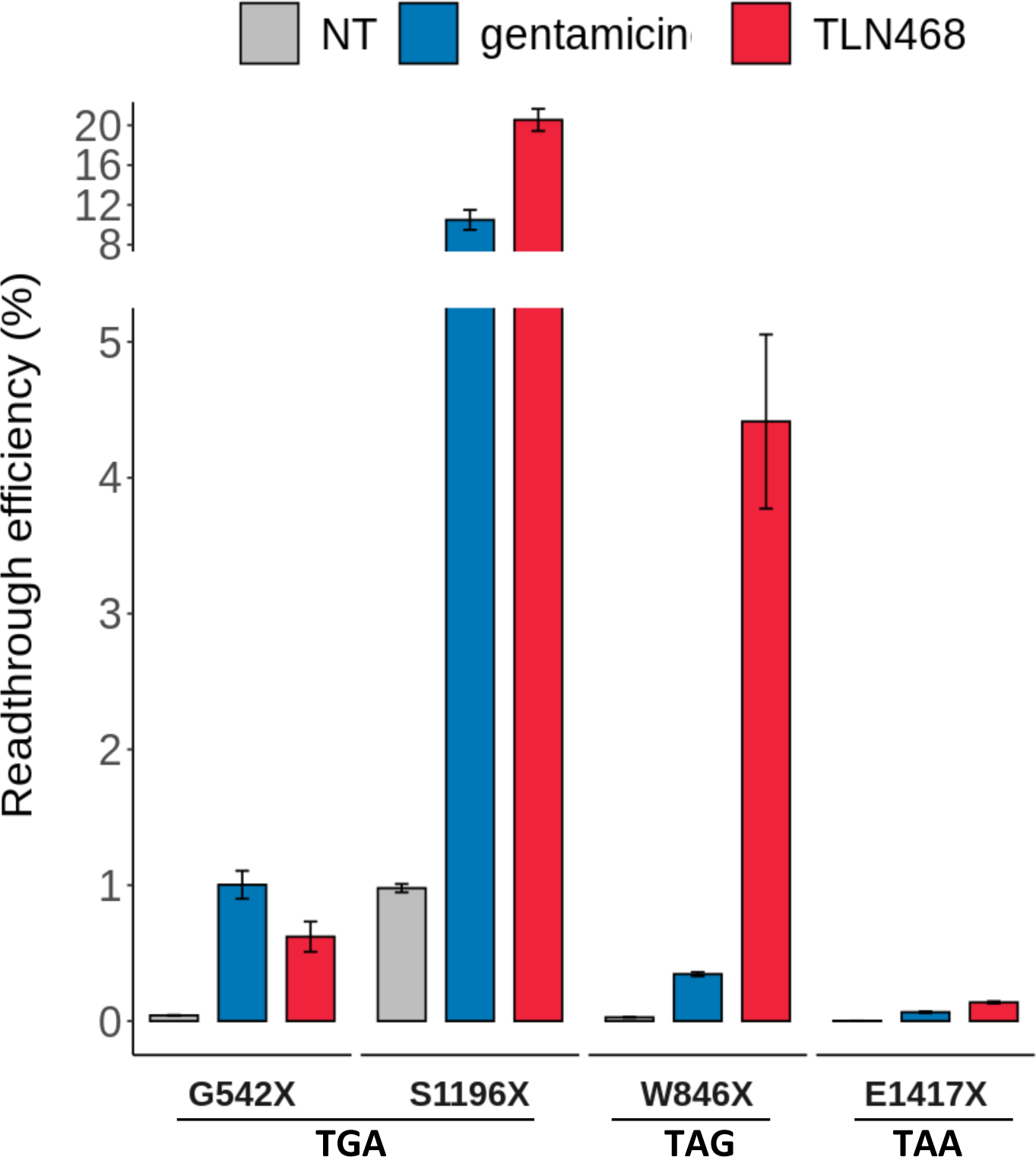
Readthrough levels in Expi293F human cells. Four CFTR PTCs (S1196X, G542X, W846X, E1417X), in basal conditions (NT) and after 24 h of incubation with gentamicin (2.5 mM) or TLN468 (80 µM) have been quantified. Bars represent the mean and standard deviation, n≥4 for all samples.

